# Demultiplexing barcoded Oxford Nanopore reads with deep convolutional neural networks

**DOI:** 10.1101/366526

**Authors:** Ryan R. Wick, Louise M. Judd, Kathryn E. Holt

## Abstract

Multiplexing, the simultaneous sequencing of multiple barcoded DNA samples on a single flow cell, has made Oxford Nanopore sequencing cost-effective for small genomes. However, it depends on the ability to sort the resulting sequencing reads by barcode, and current demultiplexing tools fail to classify many reads. Here we present Deepbinner, a tool for Oxford Nanopore demultiplexing that uses a deep neural network to classify reads based on the raw electrical read signal. This ‘signal-space’ approach allows for greater accuracy than existing ‘base-space’ tools (Albacore and Porechop) for which signals must first be converted to DNA base calls, itself a complex problem that can introduce noise into the barcode sequence. To assess Deepbinner and existing tools, we performed multiplex sequencing on 12 amplicons chosen for their distinguishability. This allowed us to establish a ground truth classification for each read based on internal sequence alone. Deepbinner had the lowest rate of unclassified reads (7.8%) and the highest demultiplexing precision (98.5% of classified reads were correctly assigned). It can be used alone (to maximise the number of classified reads) or in conjunction with other demultiplexers (to maximise precision and minimise false positive classifications). We also found cross-sample chimeric reads (0.3%) and evidence of barcode switching (0.3%) in our dataset, which likely arise during library preparation and may be detrimental for quantitative studies that use multiplexing. Deepbinner is open source (GPLv3) and available at https://github.com/rrwick/Deepbinner.

## Introduction

### Oxford Nanopore barcoding

Multiplexing (barcoding) is a common strategy used to distribute high-throughput DNA sequencing capacity over multiple samples [1]. For each input DNA sample, a unique barcode is incorporated into the library of DNA molecules prepared for sequencing. Multiple barcoded DNA libraries can then be combined and sequenced simultaneously on the same flow cell. The resulting reads must then be demultiplexed: sorted into bins according to the barcode sequence. Barcoding has obvious economic advantages, allowing users to divide the fixed cost of a sequencer flow cell over multiple input samples.

When the Oxford Nanopore Technologies (ONT) MinION sequencer was first released, its yield was measured in hundreds of Megabases (Mbp) and effective sequencing of a bacterial genome required an entire flow cell [2]. The last four years have seen a nearly 100-fold increase in yield, with 10 Gbp or more now possible from a MinION sequencing run [3]. ONT’s native barcoding kit for 1D ligation sequencing (EXP-NBD103) provides 12 barcodes which are ligated onto both ends of the DNA molecules to be sequenced. This kit allows the sequencing capacity of a single MinION run to be distributed across 12 bacterial genomes which can thus be simultaneously sequenced on a single flow cell [4].

Each ONT sequencing read is generated as a signal composed of variations in electrical current as the DNA molecule moves through the nanopore. The MinION sequencer measures the current at 4 kHz and the DNA advances at a rate of 450 bases/sec, equating to ∼9 measurements per base, on average. These ‘signal-space’ reads (a.k.a. raw signal) are translated into ‘base-space’ nucleotide sequences by basecalling software [5–7]. Basecalling is an inexact process and the resulting reads have a per-base error rate of 5–25% [8]. This error rate can be a problem for downstream analyses, including current ONT barcode demultiplexing tools, such as Albacore and Porechop, where it is common for more than 20% of the barcoded reads to be unassigned to a bin and therefore unusable [4]. Other types of ONT read analyses often achieve better performance by working with the raw signal instead [9, 10].

### Convolutional neural networks

In the last decade, neural networks – specifically convolutional neural networks (CNNs) – have revolutionised the field of image classification, achieving record high accuracies for detecting and localising objects within images [11, 12]. This progress has been fuelled by general-purpose computing on graphics processing units (GPUs) which allow much faster performance when training and classifying and have in turn allowed for more complex CNNs than were previously feasible. While early CNNs used fewer than 10 layers of neurons [13], modern varieties can be very ‘deep’, containing 100 layers or more [14]. Despite their impressive accuracy, deep CNNs have been criticised for their incomprehensibility – it can be difficult to tell how or why a CNN classifier made a particular decision [15].

Barcode classification using ONT raw signal is conceptually similar to image classification, but it is a simpler problem in two key aspects. First, ONT raw signal is a one-dimensional array of values whereas images typically have three dimensions (height, width and colour channels). Second, there are a smaller number of possible barcode classes (12 to 96, depending on the kit used) than possible image classes (often more than 1000) [16].

### Deepbinner

Here we present Deepbinner, a tool for ONT barcode demultiplexing using a deep CNN to classify reads into barcode bins using the raw read signal. We compare its performance with that of other ONT demultiplexing tools, Albacore and Porechop, which work in base-space. Operating in signal-space gives Deepbinner more power to demultiplex reads and the ability to sort raw reads for downstream uses such as Nanopolish [9]. We demonstrate its use with the EXP-NBD103 set of 12 barcodes, but it could equally be trained on any barcode set.

### Design and implementation

#### Deepbinner

##### Network architecture

Deepbinner is implemented using the TensorFlow [17] and Keras [18] code libraries. Its neural network architecture was based on elements developed in the field of image classification: groups of convolutional layers followed by max pooling layers [19]; parallel ‘inception’ modules and low dimension bottlenecks [20]; noise, dropout and batch normalisation layers [21]; and global average pooling [22].

Using these elements, we trialled hundreds of randomised network architectures to search for an effective design. Networks were assessed on their loss (categorical cross-entropy) and classification accuracy on a validation set. To discourage overfitting, we preferred models with fewer parameters and a small ratio of validation set loss to training set loss. The best performing architecture was subsequently refined to produce the final Deepbinner network shown in Fig 1. One notable way that Deepbinner’s architecture differs from image classification networks is the number of filters. Deepbinner uses a constant filter size (48, except for where the parallel module increases the filter count) whereas image classification networks commonly use a smaller number of filters in early layers and a larger number in later layers.

**Fig 1.**
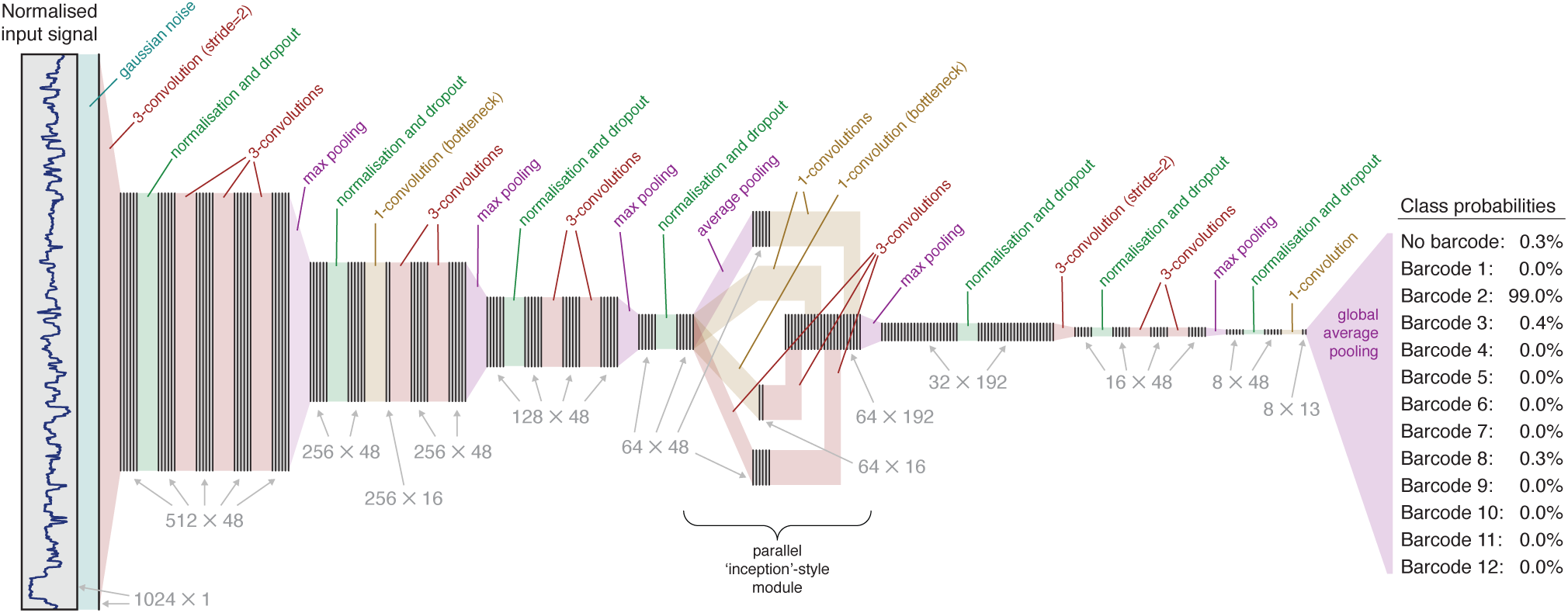
Neural network architecture. Layers in the network are drawn as coloured blocks and data as groups of vertical lines. Data dimensions are shown for each step of the process as *data length* × *filter count*. Gaussian noise and dropout layers are only active during network training, not during classification.

Deepbinner’s input array size of 1024 was chosen for two reasons. First, it is a power of two, allowing the data to halve in length as it progresses through the network’s dimension-reducing layers. Second, an ONT native barcode is 40 bp in length (24 bp for the barcode itself plus 8 bp of flanking sequence on each side) which has a typical raw signal length of 250 to 600 values, making 1024 the smallest power of two which can reliably capture the entire signal.

##### Network training

Our training data came from eight R9.4 flow cells (designed for 1D sequencing) and six R9.5 flow cells (designed for 1D^2^ sequencing but also compatible with 1D sequencing), all used with the EXP-NBD103 barcoding kit. We basecalled the reads using Albacore (v2.3.1) and searched for adapters and barcodes in the read sequences using the Edlib library [23]. Both start-read signal (raw signal from the beginning of the read) and end-read signal (raw signal from the end of the read) were extracted from each fast5 file and any open-pore signals (high current values corresponding to the absence of a DNA molecule in the nanopore [5]) were trimmed off. We then conducted semi-global dynamic time warping [24] between read signals and the expected signals for adapters and barcodes to find those elements in the read signals. Instances with a clear barcode signal were used to generate training samples of length 1024 (to match Deepbinner’s input array size).

In addition to training Deepbinner on barcoded signals, we also included a variety of barcode-less signals in the training set that were assigned a corresponding no-barcode class. These included signals from real sequences that lacked barcodes (taken from non-barcode parts of the reads), adapter signals from non-barcoded sequencing runs and multiple types of simulated signals: flat signal, Gaussian noise and Perlin noise (a coherent noise function which generates a smoother signal than Gaussian noise [25]) (Fig S1). Their presence in the training set ensures that Deepbinner can actively assign reads to a no-barcode class, not just fail to find a strong match. These no-barcode samples were included at a rate to make up approximately 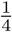 of the training samples, resulting in a total of 3 308 789 read start samples and 1 029 684 read end samples, each an array of 1024 electrical current values with a corresponding barcode label. There were fewer read-end training samples due to the fact that barcodes more reliably occur at the starts of reads than the ends.

Data augmentation is a method of artificially expanding a training set by duplicating samples with transformations [26]. Deepbinner applies data augmentation during network training by distorting signals in the temporal dimension (Fig S2). This is carried out by duplicating current values at 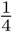 of the signal’s positions (randomly chosen) and deleting values at 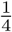 of the positions. The result is a signal of equal length but elongated in some places and shortened in others. We did not implement distortion of the signal amplitude because a similar role is carried out by the network’s Gaussian noise layer (Fig 1). For our dataset, we found that an augmentation factor of two (one augmented signal for each unmodified signal) resulted in the best classification accuracy. However, this factor is adjustable, and datasets with fewer training samples may benefit from more data augmentation.

We refined our training datasets by conducting a four-way split, classifying each quarter of the data using a model trained on the other three quarters. Samples which were misclassified (assigned a label which disagreed with the training label) were discarded. This served to remove ambiguous training samples (approximately 0.25%), leaving final training sets of 3 300 075 and 1 027 469 samples for the starts and ends of reads, respectively (totalling 16.5 GB in size).

We produced the final Deepbinner trained models by training the network for 1000 epochs (100 000 training samples per epoch) using a data augmentation factor of two and a random 95:5 training:validation split (Fig S3). Data augmentation was only performed on the training data which, combined with the network’s Gaussian noise and dropout layers (only active during training), explains why validation loss and accuracy were superior to training loss and accuracy. Each model was trained on a single NVIDIA P100 GPU and took approximately 40 hours to complete.

##### Read classification

When classifying raw ONT signal, Deepbinner will generate a probability for each possible barcode, along with a no-barcode probability (Fig 1). Deepbinner will assign a barcode to the signal if the highest probability is sufficiently larger (a difference of *>*0.5) than the second-highest probability. If the best and second-best matches are too close or if the best-match is ‘no barcode’, then Deepbinner assigns the ‘none’ label to the signal.

Deepbinner can classify each read using the start-read signal, the end-read signal or both. Using both start and end is appropriate for library preparations which add barcodes to both sides of a read, such as EXP-NBD103. If both are used, Deepbinner will independently perform classification using the read start and read end. Reads will be binned if a sufficient match was found on either end, but if the two ends match different barcodes, the read will be considered a chimera and put in the ‘none’ bin. These reads can be identified by running Deepbinner in verbose mode which provides detailed information on barcode calls. Deepbinner can optionally require a positive match on both the start and end to bin a read, enabling stringent demultiplexing. Other library preparations, such as the SQK-RBK004 rapid barcoding kit, only add a barcode to the start of the read, in which case Deepbinner’s start-only classification is appropriate.

An ONT barcode signal often appears in the first 1024 values of a read’s raw signal, but this may not be the case for reads which contain a longer-than-normal amount of open-pore signal. Deepbinner therefore examines multiple 1024-length signal windows overlapping by 512 samples. By default, it examines 11 such windows (covering 6144 samples in total), but this can be configured. Deepbinner merges the results across these windows by retaining the maximum probability for each barcode, then renormalising the probabilities.

### Evaluation

#### Study design

In order to assess the accuracy with which Deepbinner can assign reads to bins based on barcode signals, we aimed to sequence a library of barcoded DNA molecules for which the input source of each molecule could be verified independently of the barcode. To do this, we chose 12 bacterial isolates of different species that had been sequenced via Illumina HiSeq, which produces high accuracy short reads. To identify a region from each that was entirely unique to that genome, we performed a co-assembly of the pooled Illumina reads for all samples with SPAdes (v3.11.1) [27] using a small k-mer (k=23). The longest contigs from this assembly were matched to their source genome by comparison to individual (non-pooled) genome assemblies, and we choose a single contig per source genome. This produced 12 sequences, one from each genome, each composed entirely of unique 23-mers, making them easily distinguishable from each other.

#### Amplicon library preparation and MinION sequencing

The 12 bacterial isolates were grown overnight at 37°C on LB agar plates. Single colonies were then grown overnight at 37°C in Luria broth. Bacterial cell pellets from 1.5 ml of broth culture were generated by centrifugation at 15 000× g for 5 minutes. DNA was extracted from these pellets using Agencourt GenFind v2 (Beckman Coulter) with minor modifications as follows. Cell pellets were resuspended in 400 µl lysis buffer containing 9 µl Proteinase K (96 mg/ml Beckman Coulter) and 1 µl RNase A (100 mg/ml Sigma Aldrich R6513) by gentle tip mixing. Samples were lysed at 37°C for 30 minutes. gDNA was extracted from the lysed samples by completing the remaining steps of the GenFind v2 for 200 µl of blood/serum from the binding step onwards.

We used the Primer3web tool (v4.1.0) to choose 25 bp long-range PCR primers from each of the 12 unique sequences to define amplicons which ranged from 9–11 kbp (S4 File). PCR was performed using LongAMP *Taq* 2X Master Mix (New England Biolabs) with 150–450 ng gDNA as input template, primers at 1 µM, an annealing temperature of 56°C and 35 cycles of amplification. Following the PCR, size selection purification was performed with Agencourt AMPure beads (Beckman Coulter) at 0.6× ratio. The size and specificity of the PCR product was confirmed by capillary electrophoresis (Fragment Analyser AATI). A sequencing library was prepared from the purified amplicons using the Nanopore 1D ligation sequencing kit (SQK-LSK108) with the native barcoding expansion kit (EXP-NBD103) as per the manufacturer’s instructions. The run was performed on a MinION MK1b device using an R9.4 flow cell (FLO-MIN106), MinKNOW v18.03.1 and the `NC 48Hr Sequencing Run FLO-MIN106 SQK-LSK108 protocol.`

#### Demultiplexing

The reads were basecalled with Albacore (v2.3.1) using the following options: barcoding (to enable demultiplexing) and `disable filtering` (to include low-quality reads). The resulting FASTQ files were pooled and shuffled, and then given to Porechop (v0.2.3) to be independently demultiplexed. The pre-basecalled fast5 files were demultiplexed with Deepbinner (v0.2.0).

In addition to running the tools with default parameters, we also tested Porechop and Deepbinner with parameters to increase or decrease their demultiplexing stringency. Porechop’s lenient settings reduce the barcode and difference thresholds from their default values (75% and 5%, respectively) to 60% and 1%. This makes Porechop willing to consider very low quality alignments and assign a barcode even when there is a close second-best match. Porechop’s stringent settings increase the barcode threshold to 85% (so it will only consider high quality alignments) and require a barcode match on both the start and end of reads. The only difference between Deepbinner’s default and stringent settings is that the latter requires a barcode match on both the start and end of reads.

We assigned ground truth classifications to the ONT reads by aligning their basecalled sequences to the amplicon reference sequences with minimap2 (v2.12) [28] and binning with the `assign reads to reference.py` script (included with Deepbinner). Reads which failed to meet an alignment threshold (100 bp or 10% of the read length, whichever is smaller) were classified as ‘unknown’. Reads which exceeded an alignment threshold to a secondary amplicon (50 bp or 5% of the read length, whichever is larger) were classified as ‘chimera’. This method is only able to detect cross-bin chimeras, i.e. reads with separate components from two different amplicons. It cannot detect within-bin chimeras, e.g. two copies of the same amplicon concatenated in a single read.

#### Additional test sets

We also performed a whole genome sequencing (WGS) run using 12 different bacterial species, as it represents a more realistic sequencing scenario for many users. We followed the same preparation and analysis described above for the amplicon sequencing but with genomic DNA and excluding the PCR-specific steps. The reads were aligned with minimap2 (v2.12) [28] to all twelve complete genome sequences (produced via hybrid Illumina-Nanopore assembly using Unicycler v0.4.6 [4]) and assigned ground truth classifications with the same script and thresholds used for binning the amplicon reads. Due to shared sequences between the genomes, confidently assigning ground truth was more challenging than for the amplicon set, resulting in more reads being classified as ‘unknown’.

Based on examination of basecalled sequences, we estimate approximately 0–2% of reads in a barcoded read set may be genuinely lacking a barcode, i.e. one was not ligated during preparation. These reads can be difficult to distinguish from reads which do have a barcode but where no classification was possible, so to examine the demultiplexing tools’ behaviour on non-barcoded reads, we additionally assessed them using a non-barcoded ONT read set (SQK-LSK108, R9.4 flow cell). This is referred to as the ‘negative control’ set and the ground truth classification for all its reads was ‘none’.

To ensure a fair assessment, no reads from any of the three test sets (amplicon, WGS and negative control) were used in the training of Deepbinner’s models.

## Results

### Performance on test sets

The barcoded amplicon MinION sequencing run produced 1 893 881 reads (9.69 Gbp), 1 642 052 of which (87%) could be reliably assigned to an amplicon based on the internal read sequence alone, producing our ground truth classifications. The WGS run produced 1 300 656 reads (12.2 Gbp), 968 137 of which (74%) could be assigned to a genome for ground truth. Our negative control set contained 101 993 reads (945 Mbp).

Of all three demultiplexers tested (using default parameters), Deepbinner performed best on both precision (a.k.a. positive predictive value, proportion of binned reads correctly assigned) and recall (a.k.a. accuracy, proportion of all reads correctly assigned) (Table 1, S1 File). It was particularly strong on recall, correctly binning 253 300 more reads (1265 Mbp) than Albacore and 176 740 (836 Mbp) more than Porechop in the amplicon set. The same patterns were evident on the WGS set, with Deepbinner displaying improvements of ∼10% in recall and 1–2% in precision, as compared to Albacore and Porechop (S2 File).

**Table 1.**
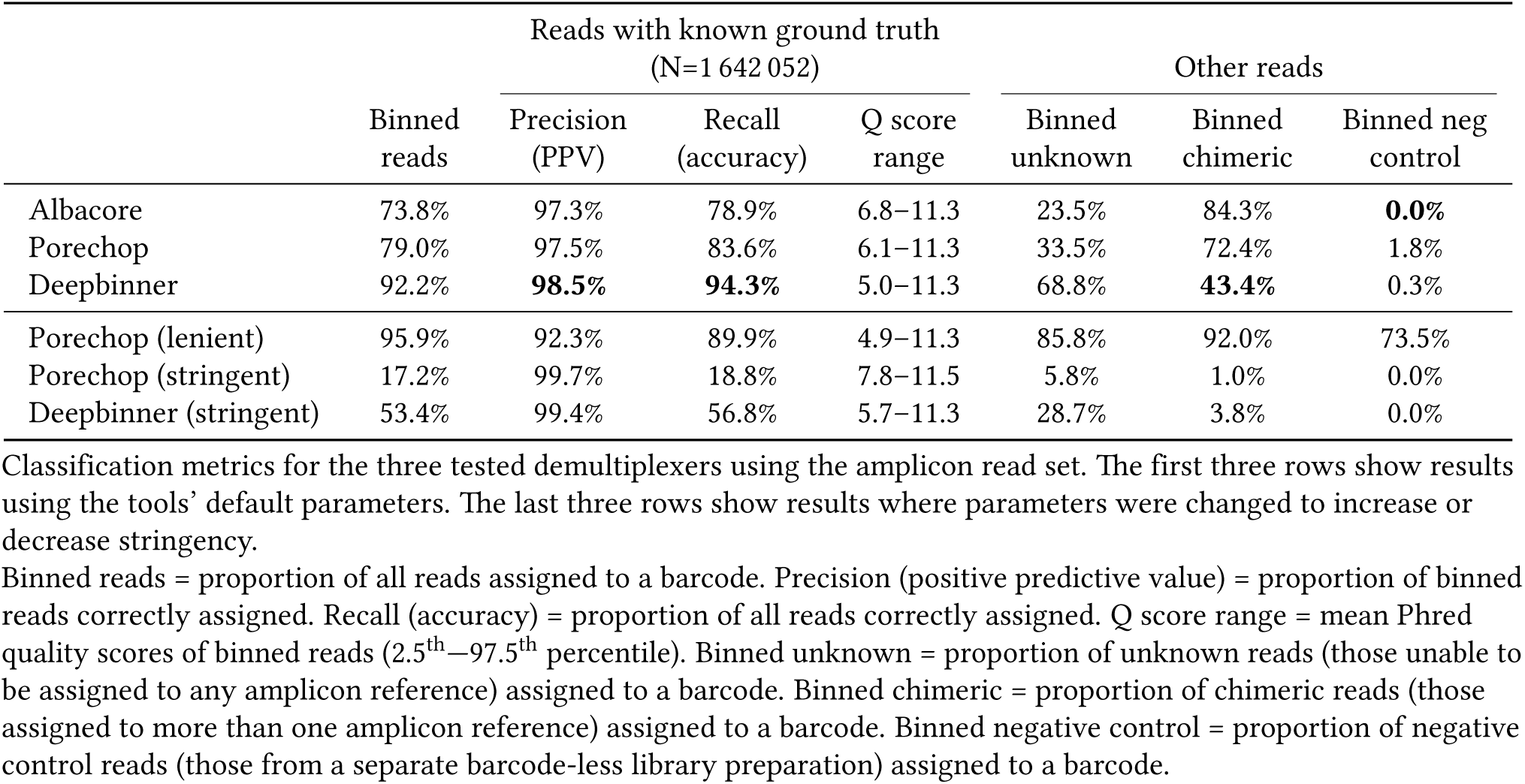
Classification performance of demultiplexing tools.

Deepbinner’s superior recall is primarily due to its ability to bin low quality reads. All three demultiplexers perform similarly on very high quality reads, but Albacore and Porechop’s recall fall below 50% for low quality reads (Fig S4). Deepbinner, however, maintains a recall of over 85% for the entire quality range.

Deepbinner was much more likely than other demultiplexers to put ‘unknown’ reads (those which we could not assign to a reference sequence, 12.7% of the amplicon reads) into a barcode bin. Some are likely reads that were too low quality to be assigned to a source based on the internal sequence. The negative control test shows that both Porechop and Deepbinner can suffer from over-sensitivity, putting barcode-less reads in a bin. For chimeric reads (0.31% of the amplicon reads), Deepbinner was less likely than other tools to assign a barcode.

There is often a trade-off between precision and recall which can be adjusted via demultiplexer parameters (Table 1). When Porechop is given lenient settings, it has lower precision (92.3%) but improved recall (89.9%). When given stringent settings, Porechop achieves very high precision (99.7%) but at great cost to recall (18.9%). When Deepbinner is given stringent settings, its precision and recall become 99.4% and 56.8%, respectively. Combining multiple demultiplexers (only binning reads where multiple tools agree) improves precision at the cost of recall (S1 File). However, precision never exceeded 99.7%, even when we used multiple demultiplexers and very stringent settings.

Deepbinner requires as little as 1 CPU and 1 GB of RAM, though it can take advantage of additional resource to run faster, benefitting from up to about 16 CPUs and 8 GB of RAM. It is difficult to make a speed performance comparison between Deepbinner and the other tools, as they use computer resources differently. Albacore is likely the fastest for many users, as its demultiplexing requires little extra time over basecalling. If Deepbinner is run on a CPU, it is the slowest demultiplexer tested – classifying the amplicon set took 29.5 hours to complete (∼18 reads/sec, using a total of 347 CPU hours). When run on a GPU (NVIDIA P100), its performance is comparable to Porechop, with the amplicon set classification taking 4.6 hours (∼114 reads/sec).

### Implications

Many of Deepbinner’s advantages stem from the fact that it operates not on basecalled sequences, but on the more informative raw signal. This may account for Deepbinner’s high recall (including the ability to bin lower quality reads) and precision. By demultiplexing fast5 files before basecalling, it simplifies downstream analyses such as Nanopolish which require raw reads [9]. Deepbinner also opens up the possibility of constructing barcodes with modified DNA bases to increase the size of the genomic alphabet. Such high-alphabet barcodes could be easier to differentiate at the signal level, and if so would allow for a greater number of unique barcodes in a given sequence length.

The disadvantages of Deepbinner are similar to those experienced by CNN classifiers in other contexts. Training the network is computationally intensive, and a large volume of training data is required. Trained networks may not generalise well across different flow cells and library preparation kits, necessitating a separate trained network for each. When Deepbinner makes an error during classification, the ‘black box’ nature of neural networks makes it difficult to understand why.

### Recommendations

Deepbinner’s high recall is well suited to applications where the most important factor is maximising the number of classified reads and therefore the amount of useable data. A user can run Deepbinner during sequencing, binning fast5 files as they are produced, and then run Albacore on each resulting directory of reads. While other demultiplexing approaches can discard (i.e. fail to classify) over 20% of the data, this proportion will likely be less than 10% with Deepbinner. This greater yield of reads may improve assemblies, even when the additional reads are low-quality [29].

For applications where precision is paramount (i.e. incorrectly binned reads must be minimised), Porechop with stringent settings may be the best choice, though its very low recall means that most reads (*>*80%) will be lost. Deepbinner with stringent settings (requiring both start and end barcodes) has slightly worse precision than stringent Porechop but can classify more than half the reads.

Our tests showed that demultiplexing precision plateaued at about 99.7%, i.e. 0.3% of reads with a ground truth label are consistently assigned to the wrong bin by all demultiplexing tools. This implies these reads may have the wrong barcode ligated to the DNA, a problem that no demultiplexing tool could fix. This may arise in the library preparation, whereby unligated barcodes could be carried through after sample pooling and then be available for ligation to incorrect DNA fragments in the adapter ligation step. If so, a bead clean-up with size selection after barcode ligation (but before adapter ligation) may mitigate the issue by reducing the number of free barcode sequences. This small amount of barcode crosstalk, along with cross-barcode chimeric reads, is likely inconsequential for isolate sequencing but could be a serious problem in some quantitative applications such as metagenomics or transcriptomics [30].

## Availability and future directions

Deepbinner, documentation and pre-trained models are available on GitHub: https://github.com/rrwick/Deepbinner. The raw fast5 read sets used in this manuscript are available on ENA: https://www.ebi.ac.uk/ena/data/view/PRJEB28450. The basecalled reads and full classification results are available on figshare: https://figshare.com/projects/Deepbinner/34223.

Future development of Deepbinner will involve training models for other library preparations, including larger barcode sets (more than 12 barcodes) as they become available. Improved performance and parallelism is also a focus, to ensure that Deepbinner can keep up with high yield sequencing runs, such as those from ONT’s higher throughput GridION X5 and PromethION sequencers.

## Acknowledgments

We would like to thank Wendy Wilson for conducting the long-range PCR amplification.

## Supporting information

**S1 File.** Confusion matrices (amplicon). Classification counts per reference bin and predicted bin, for each demultiplexer tested, using the amplicon read set.

**S2 File.** Confusion matrices (WGS). Classification counts per reference bin and predicted bin, for each demultiplexer tested, using the whole genome read set.

**S3 File.** Negative control results. Classification counts per predicted bin, for each demultiplexer tested, using the negative control read set.

**S4 File.** Amplicon sequences. FASTA file of the 12 amplicon sequences.

**Fig S1.**
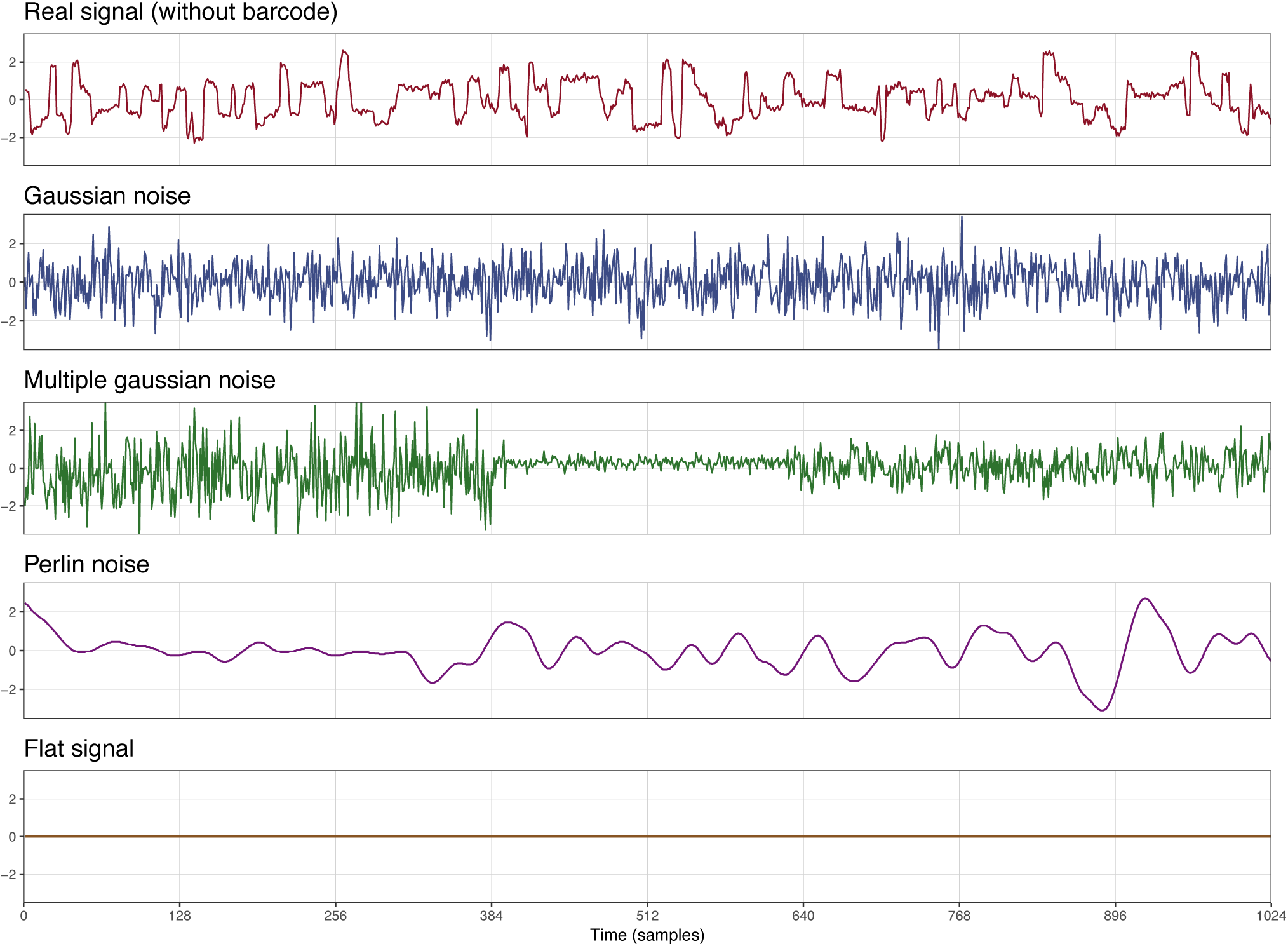
No-barcode training signals. Multiple types of signals were included in the training set to explicitly teach the neural network what a barcode-free signal looks like. The signal amplitude has been normalised to a mean of 0 and a variance of 1.

**Fig S2.**
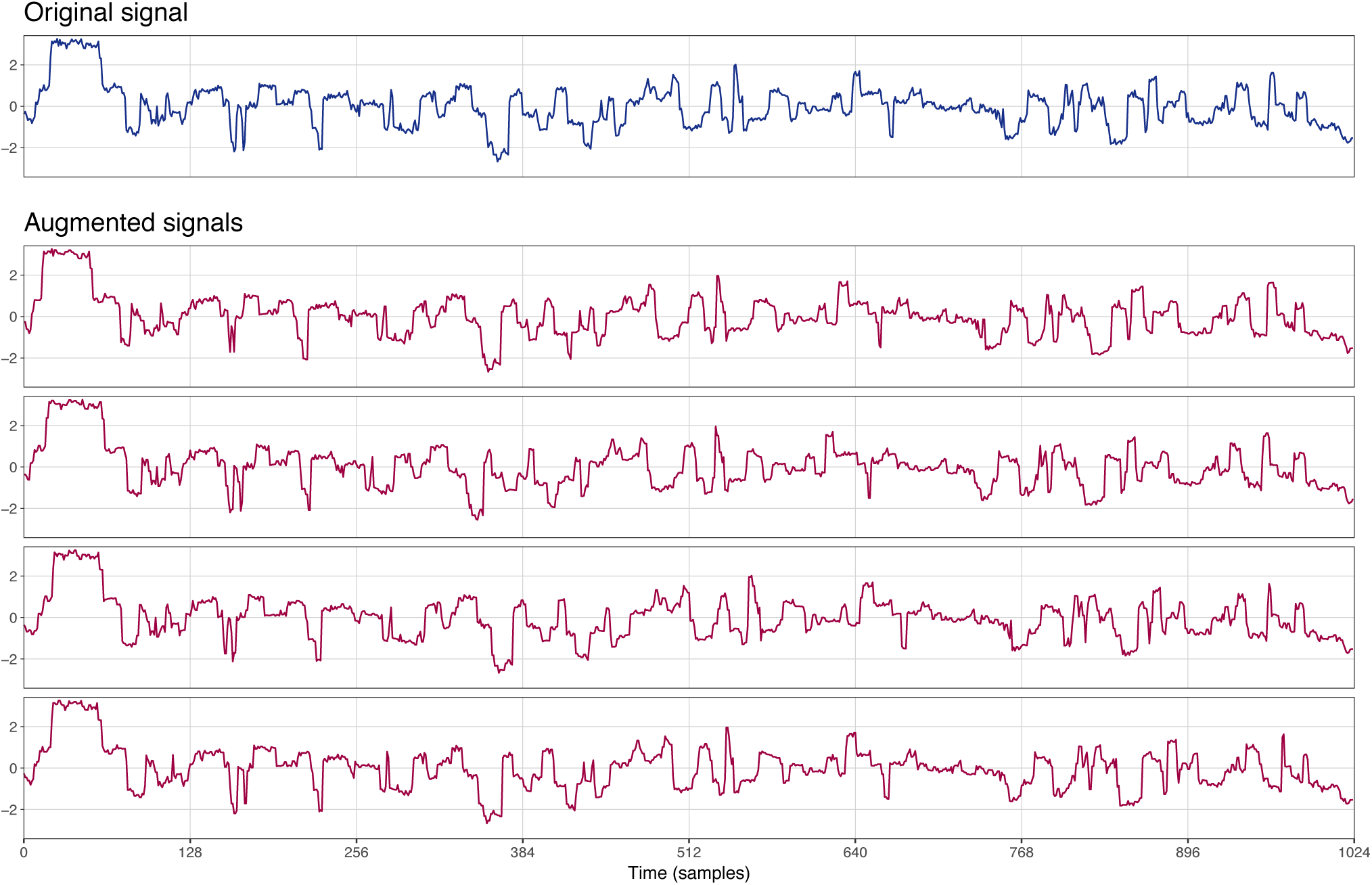
Data augmentation via temporal distortion. One real training sample can yield multiple additional training samples by distorting the signal along the temporal axis. The signal amplitude has been normalised to a mean of 0 and a variance of 1.

**Fig S3.**
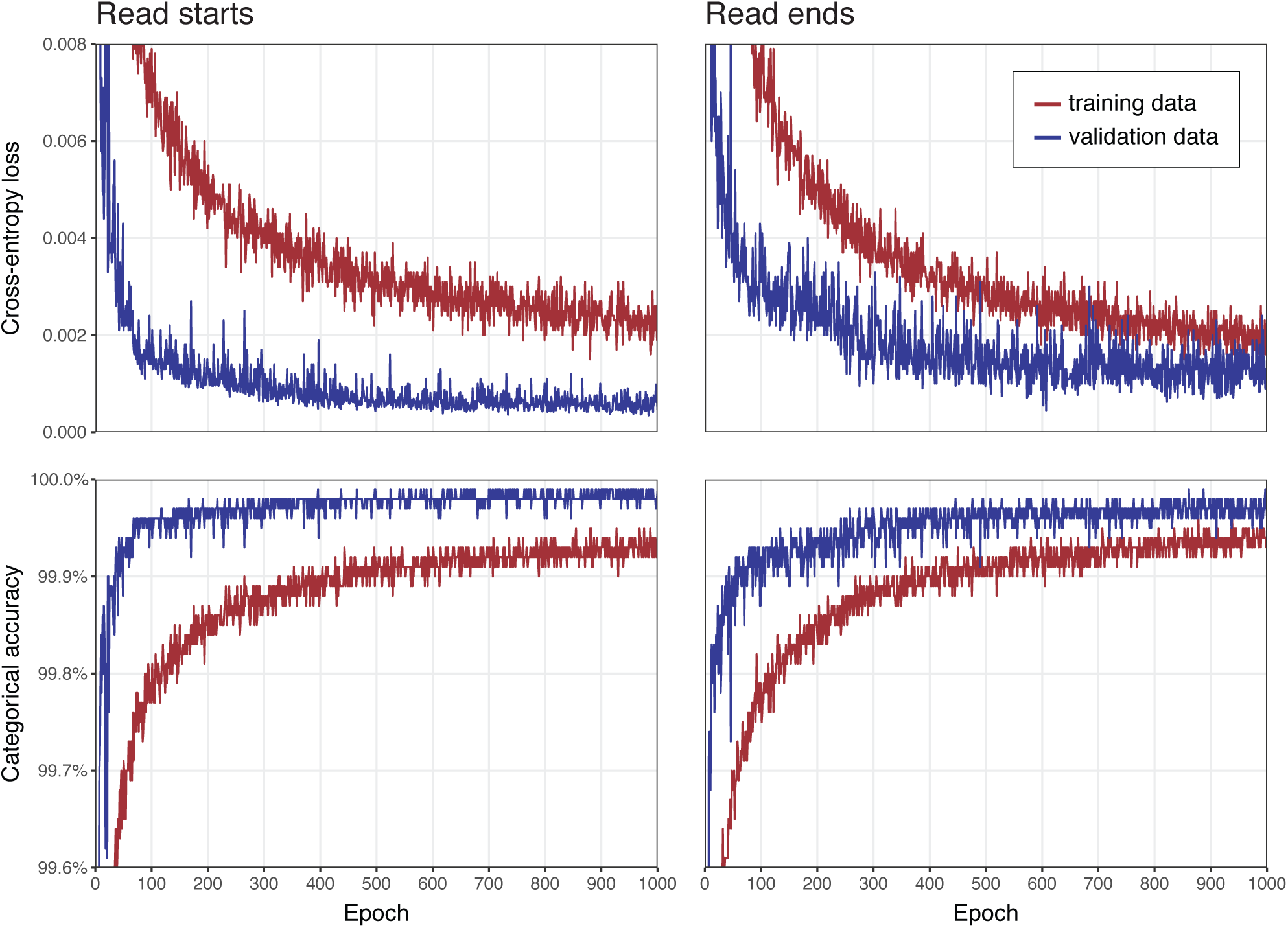
Training metrics for the read start and read end models. Generated using a random 95:5 training:validation split. Training data has poorer performance than validation data due to data augmentation and training-only layers (Gaussian noise and dropout) in the network.

**Fig S4.**
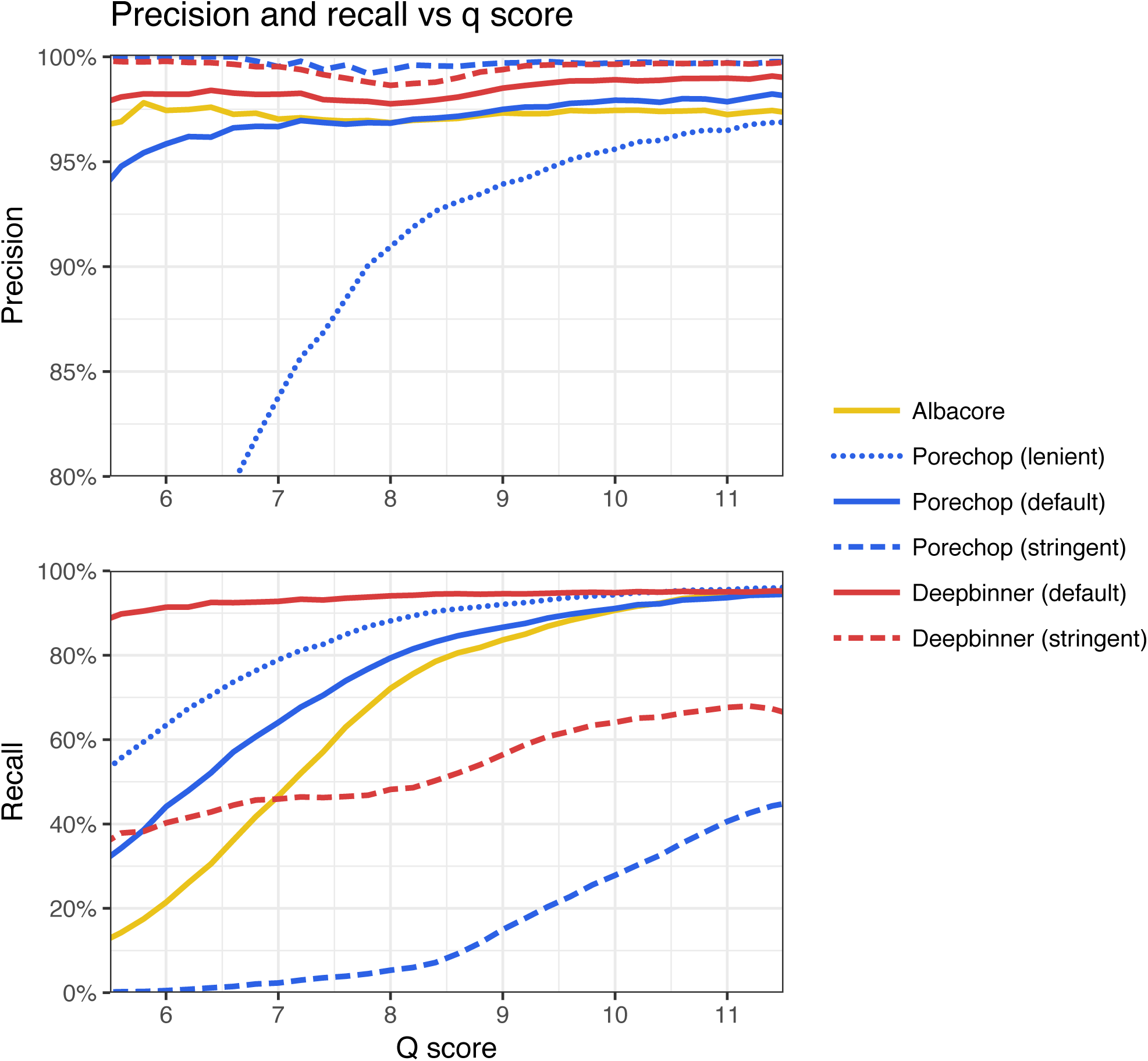
Classification metrics by q score. Precision and recall for each tool’s demultiplexing of the amplicon read set as a function of read q score.

